# Theta oscillations shift towards optimal frequency for cognitive control

**DOI:** 10.1101/2020.08.30.273706

**Authors:** Mehdi Senoussi, Pieter Verbeke, Kobe Desender, Esther De Loof, Durk Talsma, Tom Verguts

## Abstract

Cognitive control is supported by theta band (4-7Hz) neural oscillations coordinating neural populations for task implementation. Task performance has been shown to depend on theta amplitude but a second critical aspect of theta oscillations, its peak frequency, has mostly been overlooked. Using modelling, behavioral and electrophysiological recordings, we show that theta oscillations adapt to task demands by shifting towards the optimal frequency.

---

Cognitive control permits adapting behavior to task demands, crucial in an ever-changing environment. It is supported by neural oscillations in the theta band (4-7Hz)^1^ that coordinate distant neural populations to create task-relevant functional networks through synchronization^2^. To be adaptive, theta oscillation characteristics must change with task demands. Most research has focused on theta amplitude, showing that it increases after conflicts and errors, causing subsequent neural adaptation leading to better task performance^1^.

A second aspect of theta, its peak frequency within the 4-7Hz range, also varies across tasks and participants^3^. However, this variability is commonly ignored and no theory considered its role in cognitive control. To address this gap, we draw from two prominent frameworks: biased competition (BC)^4^ and communication-through-coherence (CTC)^5^. We built a novel computational model where theta oscillations orchestrate competition between task representations, which in turn guides CTC to set up task-relevant functional networks. Model simulations show that, depending on task demands, different theta frequencies were optimal for task performance. We tested model predictions on behavioral and electrophysiological data and confirmed that theta peak frequency shifts towards optimal frequency depending on task demands.

We designed a novel stimulus-action mapping task (**Fig.1a**) wherein on each trial, a different mapping (i.e. a rule, with variable difficulty) needs to be established. The task consists of reporting the tilt of one of two gratings, clock-wise (CW) or counter-CW from the vertical axis, using the index or middle finger of one of both hands. On each trial a two-letter cue instructed the rule: which was the target grating (Left (L) or Right (R), top-letter) and which hand to use (L or R, bottom-letter). We thus manipulated task difficulty: same-side cues (i.e. (top-letter – bottom-letter) RR and LL) were easier than different-side cues (LR and RL).

**Figure 1.**
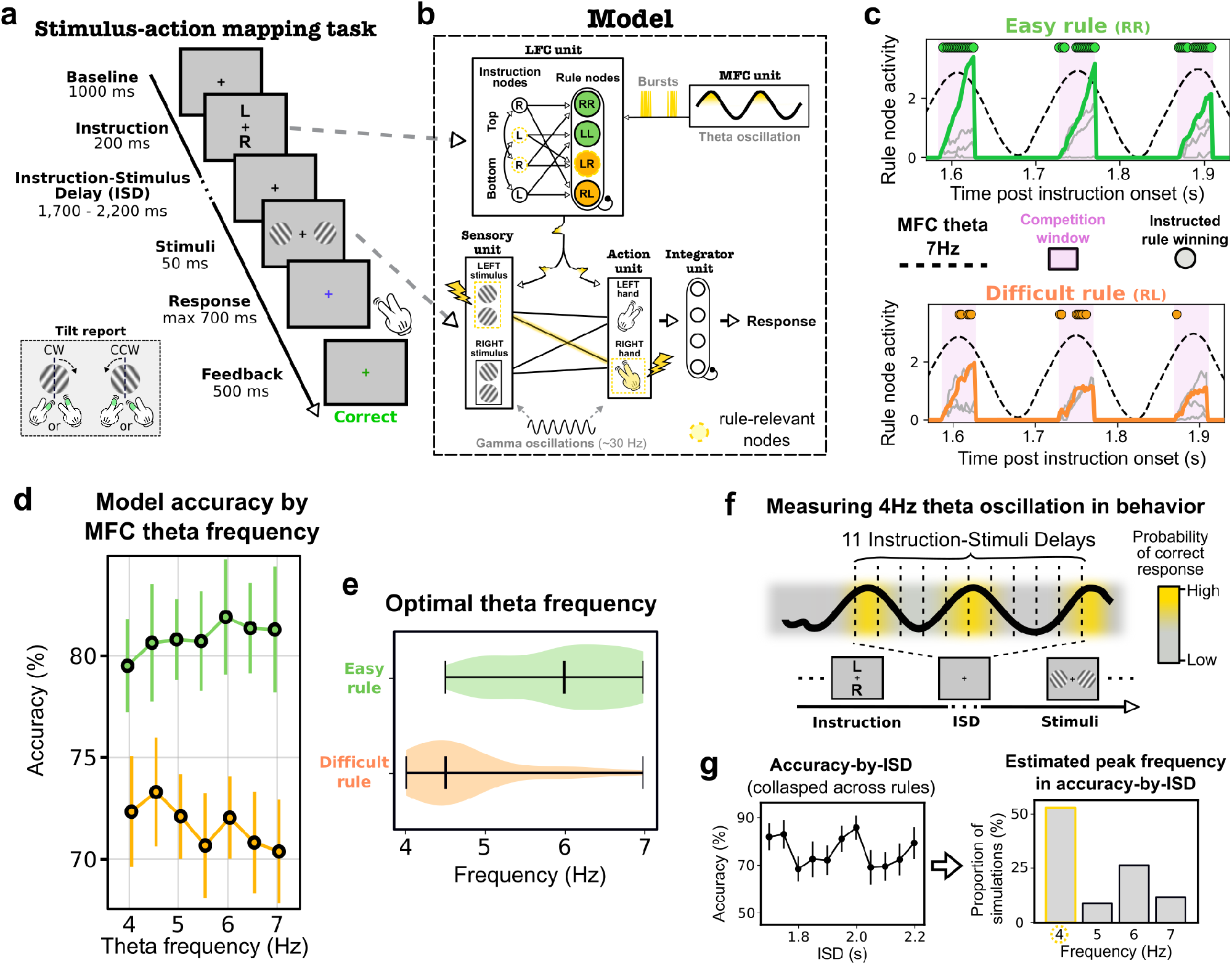
Model and simulations. **a.** Stimulus-action mapping task. Each trial starts with a cue instructing the mapping to use. In this example the rule is “LR” instructing to report the left grating’s tilt with the right hand. **b.** Model architecture. **c.** Rhythmic BC in rule nodes: 3 cycles at a fast MFC theta frequency (7Hz). Left panel (Easy rule): the rule node (green curve) rapidly wins the competition over other rule nodes (grey curves). Right panel (Difficult rule): the rule node (orange curve) struggles to win the competition and often loses to other rule nodes (grey curves). **d.** Model accuracy by rule difficulty across theta frequencies. **e**. Theta frequency yielding highest model accuracy per rule difficulty. **f.** Measuring theta oscillations in behavior through densely-distributed ISD. **g.** Accuracy-by-ISD (left) and estimated peak frequency (right). All error bars represent s.e.m.

Our model consists of four main units (**Fig.1b**): two control units (lateral and medial frontal cortex, LFC and MFC respectively), and two processing units (Sensory and Action). In LFC, cues activate instruction nodes, which themselves activate rule nodes. Rule nodes form a competitive accumulator network, and implements BC: In a Stroop-like manner, the connectivity between instruction nodes induces stronger competition between rule nodes for different-side than same-side rules. Importantly, rule node competition is orchestrated by theta oscillations generated by the MFC unit: competition is (re-)initiated when MFC theta exceeds a processing threshold (**Fig.1c**). Each rule node points to rule-relevant processing modules. Processing nodes oscillate at gamma frequency. Rule nodes gate communication between Sensory and Action units through CTC^5,6^ by means of phase-resetting bursts emitted by MFC at theta oscillation peaks^2,6^ (**Supplementary Fig.1c**).

Critically, with a fast theta frequency, e.g. 7Hz, rule nodes gate processing modules frequently, shortening “off”-periods in which rule-relevant processing nodes de-synchronize, at the cost of shorter competition windows. With a slow theta frequency, e.g. 4Hz, gating is imposed less frequently, but competition windows are longer. Due to BC, one rule will win the competition; but for difficult rules, resolving the competition will take more time, i.e. require longer competition windows. In our task different-side rules are more difficult, so the model achieves better performance at slower theta frequencies where competition is long enough for the correct rule node to win. In contrast, for easy rules, higher theta frequencies yield better performance because the competition is won quickly and rule-relevant nodes are frequently gated. Hence, an optimal agent would shift theta frequency depending on task demands.

Model simulations (**Fig.1d**) confirmed that for difficult rules the model achieves optimal accuracy at a slow theta frequency, whereas for easy rules, a fast theta is optimal (*t*(33)=5.8, *p*<10^−5^, **Fig.1e**). Fits from the drift diffusion model (**Methods**) showed that only drift rate exhibited this theta-frequency – rule-difficulty interaction (**Supplementary Fig.5a**), refuting a speed-accuracy trade-off (SATO) explanation. Theta amplitude alone could not explain this result as it only negligibly affected competition window length as compared to frequency (**Supplementary Fig.2c**). Furthermore, theta-rhythmic gating of processing nodes should yield higher model performance shortly after a burst, i.e. at theta oscillation peaks (**Fig.1f**). By varying the instruction-stimulus delay (ISD), to sample model performance at different phases of the theta-rhythmic process, we showed that model accuracy oscillates at a frequency closely matching MFC theta frequency (*p*<10^−5^; **Fig.1g**, **Supplementary Fig.3**).

These simulations lead to two key behavioral and neural predictions. First, oscillations of accuracy-by-ISD should shift towards optimal theta frequency depending on task demands; second, frontal theta oscillations should also exhibit this effect, which should predict subsequent task performance.

In an experiment on human participants, we first confirmed that rules varied in difficulty. There was a significant target-location – hand interaction in accuracy (RR and LL easier than LR and RL; *F*(1,32)=24.29, *p*<0.00003, **Fig.2a**), and a main effect of hand (*F*(1,32)=12.69, *p*=0.00117). Consistent with model simulations, only drift rate exhibited this interaction (**Supplementary Fig.5b**); we therefore used accuracy as our dependent variable. To test model predictions on behavioral oscillations, we computed peak theta frequency of accuracy-by-ISD (**Methods**). As predicted, we found a significant target-location – hand interaction (*F*(1,32)=7.53, *p*=0.00098), showing that accuracy oscillated at a slower theta frequency for difficult rules (LR, RL; **Fig.2b**), and no main effect.

**Figure 2.**
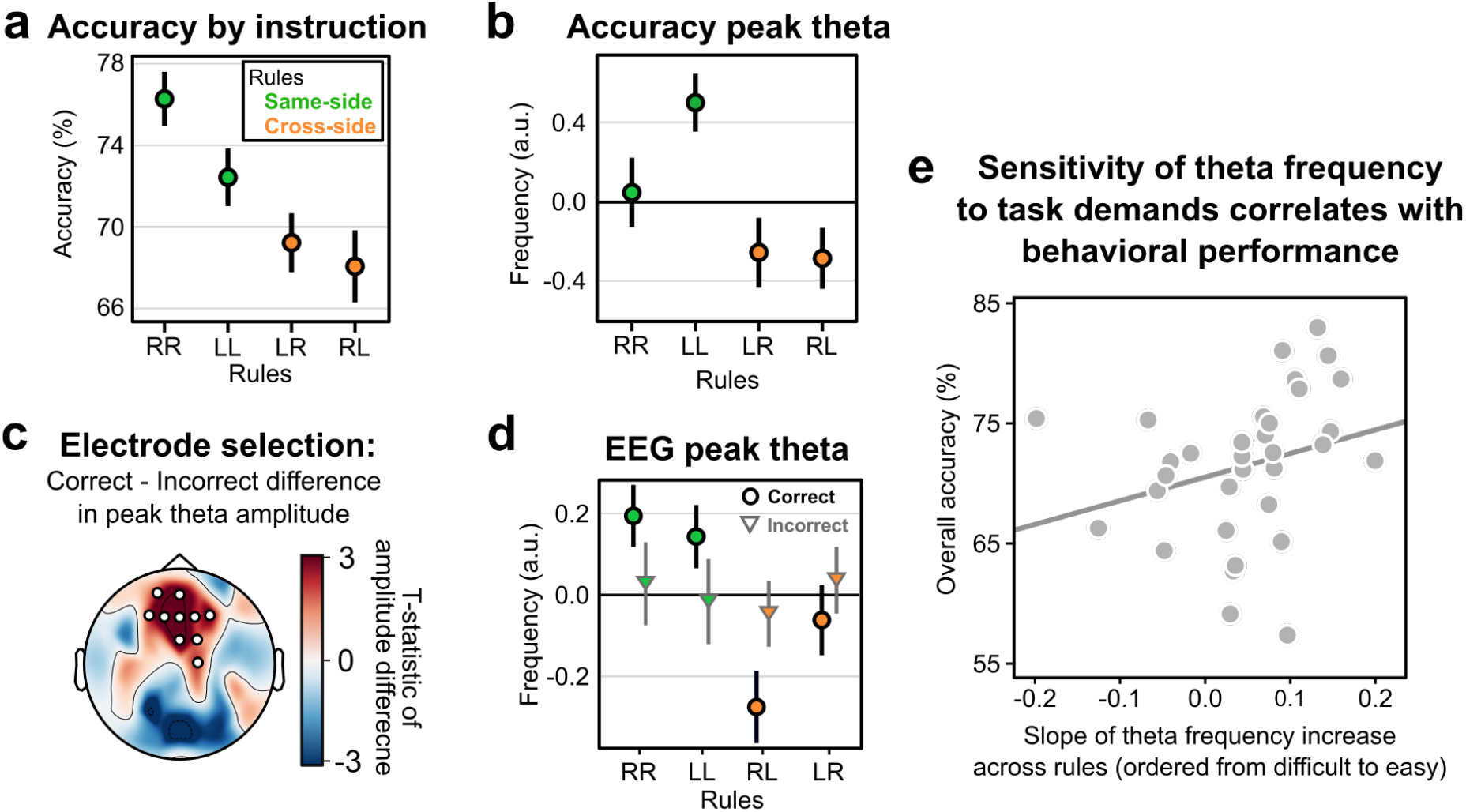
Testing model predictions in behavior and EEG. **a.** Overall accuracy. **b.** Peak frequency of oscillations in accuracy-by-ISD. **c.** Frontal cluster of electrodes with increased theta amplitude. **d.** Theta peak frequency by rule in frontal electrodes cluster. **e.** Correlation between theta peak slope across rule difficulty and overall accuracy. All error bars represent s.e.m.

Next, we investigated whether neural theta exhibited this frequency shift due to task demands. We extracted EEG theta peak *frequency* in a 1s pre-stimulus window from an electrode cluster exhibiting significantly higher theta *amplitude* in correct than incorrect trials (*p*<10^−4^; **Methods**; **Fig.2c**). As predicted, peak theta *frequency* in correct trials significantly decreased between same-side and different-side rules (*F*(1,32)=12.31, *p*=0.00136; **Fig.2d**). Furthermore, contrasting correct and incorrect trials revealed that higher theta frequency improved performance in same-side rules, whereas a lower theta frequency improved performances in different-side rules (*F*(1,32)=5.80, *p*=0.02193; **Fig.2d**). Finally, across participants, the degree to which theta frequency shifted from difficult to easy rules positively correlated with overall accuracy (*r*=0.43, *p*=0.01187; **Fig.2e**), indicating that a higher sensitivity of theta frequency to rule difficulty improved task performance. These results cannot be explained by theta amplitude alone as both peak and amplitude were estimated independently over the 1/f spectrum (**Supplementary Fig.4c**).

These findings align with evidence that frequency of neural oscillations adapt to external demands, e.g. perceptual demands in alpha band^7^, and is related to short-term memory capacity in theta band^8^.

Neural oscillations may address the fundamental binding problem in cognition by gating information flow in the brain. Our results provide critical insights into the adaptive nature of theta oscillations supporting cognitive control, and call for a more systematic evaluation of theta characteristics, at computational, behavioral, and neurophysiological levels.

## Author contribution

M.S., D.K. and T.V. designed the study. M.S., P.V. and T.V. developed the model. M.S. and E.D.L. collected the data. M.S. analyzed model simulations, behavioral and EEG data. M.S. and T.V. wrote the manuscript. All of the authors discussed the results and commented on the manuscript.

## Acknowledgments

The authors want to thank Cristian Buc Calderon for fruitful discussions and comments on the manuscript. MS and TV were supported by grant G012816 from Research Foundation Flanders. TV and EDL were supported by grant BOF17-GOA-004 from the Research Council of Ghent University. PV was supported by grant 1102519N from Research Foundation Flanders. KD was supported by the FWO [PEGASUS]^2^ Marie Skłodowska-Curie fellowship 12T9717N.

## Methods

### Model

#### Overview

The model implements biased competition (BC) and communication-through-coherence (CTC) and consists of five units: two control units (lateral and medial frontal cortex, LFC and MFC respectively), two processing units (Sensory and Action units), and an Integrator unit accumulating evidence from the Action unit, and producing a response. We will first briefly describe how BC and CTC are implemented in the model, and then proceed to a detailed description of each unit, and the nodes composing them.

BC proposes that task representations compete, biased by top-down input. We implemented BC in the LFC unit, which was composed of rule nodes that pointed to specific processing nodes. Each rule node pointed to processing modules composing the rule. This allows a rule node to gate task representations (encoded via an input-output matrix), relevant for that particular rule. For instance, a rule node could implement the rule “report sensory feature 1 using action set 2” (see this example in **Fig.1b** and **Supplementary Fig.1a**). We used location (Left (L) or Right (R)) as a sensory feature. We used two action sets, namely Left (L) and Right (R) hand (see Action unit in Fig.1b). Rule nodes were interconnected to create a competitive accumulator network. Each rule node also received a biasing input throughout a trial from instructions in the form of two letters presented simultaneously and modeled as a top letter instructing which stimulus feature was the target (L or R) and bottom letter instructing which action set to use (L or R). We refer to these instructions, or rules, in this manner: RL for “Right-Left”, in which the first letter refers to the top instruction letter, instructing the target stimulus feature (Right grating), and the second letter refer to the bottom instruction letter, instructing the action set to use (Left hand). Each rule in the task (i.e. RR, LL, LR, RL) activated a unique set of instruction nodes (see in **Fig.1b** and **Supplementary Fig.1a**, LFC unit). Two nodes represented the top letter of an instruction, and two others the bottom letter. This network of instruction nodes created a congruency effect between instruction letters: top and bottom “Left” nodes were connected, thereby activating each other, and similarly for “Right” nodes. In a Stroop-like manner, the connectivity in instruction nodes induced a stronger input to rule nodes for same-side (LL, RR) than for different-side (LR, RL) rules. Furthermore, different-side rules also activated non-instructed instruction nodes more than same-side rules due to the lateral excitation in instruction nodes, thereby making the BC between rule nodes more difficult for different-side rules to win.

The top-down bias signal from control units was implemented through CTC. The MFC unit generated theta oscillations. During a temporal window whose size depended on the specific theta frequency (i.e. the slower theta, the longer the temporal window), a competition was initiated between rule nodes. During this competition window, MFC unit sent bursts of activity^6,9,10^. The most active rule node (i.e. the one “winning” the competition) amplified the burst and sent it to Sensory and Action nodes it points to. All Sensory and Action nodes oscillated at gamma frequency. These bursts reset the phase of Sensory and Action nodes selected by the LFC unit, and increased synchrony between them, allowing for efficient communication, i.e. gating. Through this selective routing of bursts to Sensory and Action nodes, the model implements CTC by creating functional networks to implement a rule.

As a result of the BC, one rule (typically, the correct one) will win the competition; but in cases in which the competition is stronger, it will require a longer competition window for the correct rule to win the competition. The latter are difficult rules. In the model, rule difficulty was implemented through conflicting instructions that activated more rule nodes than easy instructions, making the competition more balanced between the instructed and the other rule nodes.

The consequence is that for difficult rules, the model will achieve better performance at a slower theta oscillation frequency because longer competition will permit rule nodes to win the competition and thus gate the rule-relevant processing nodes. In contrast, for easy rules, performance increases with a higher theta frequency because rule nodes quickly win the competition and the faster theta frequency allows to more frequently gate rule-relevant processing nodes. Hence, an adaptive agent would shift theta frequency depending on task demands.

**Supplementary Figure 1.**
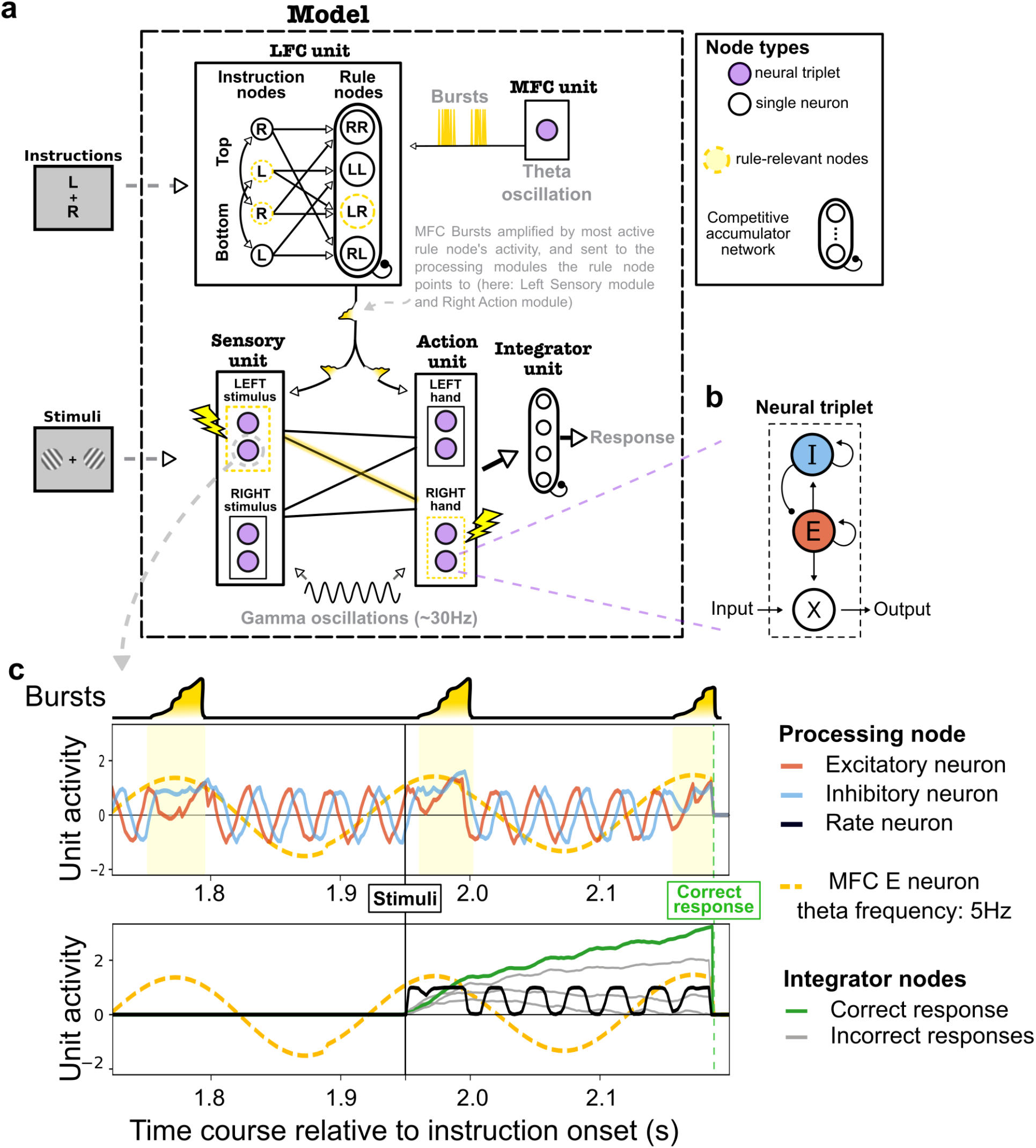
Model structure and dynamics. **a.** The model is composed of five units: two control units (LFC and MFC), two processing units (Sensory and Action unit), and an Integrator unit. On each trial instructions activated instruction nodes, biasing the competition between rule nodes which was initiated when MFC theta oscillations exceed a processing threshold. The most active rule node, during each competition window, routed bursts emitted by the MFC unit at the peak of theta oscillations to rule-relevant processing nodes, thereby phase-aligning (synchronizing) their gamma oscillations. At stimuli presentation, Sensory nodes activated by the stimuli activated Action nodes corresponding to the action associated with each grating’s tilt. Each action node activated one Integrator node. Integrator nodes constituted a competitive accumulator network. When one of the nodes reached a response threshold, a response (e.g. left-middle finger) was recorded, as well as the reaction time relative to stimuli onset. **b.** Each node of MFC and processing (Sensory and Action) units is a neural triplet node. This neural triplet is composed of one excitatory (E), one inhibitory (I), and one rate neuron (x). The E-I pair generated oscillations at frequency depending of their coupling parameter. Bursts coming from control units were sent to E neurons. Rate neurons received input from, and sent output to, other nodes’ rate neurons. The activity (output) of a rate neuron is modulated by the E neuron of its neuronal triplet. **c.** Time course of neural triplet activity in the rule-relevant Sensory unit around stimuli presentation.

#### Oscillatory nodes: a neuronal triplet

In the MFC unit and processing units, each node *i* implements a cortical column simplified as a triplet of neurons: a rate code neuron (*x_i_*) and two phase neurons (one excitatory (*E_i_*) and one inhibitory (*I_l_*) neuron), see Supplementary **Fig.1a-b**.

Phase neurons, i.e. the E-I pair, generate oscillations with a frequency defined by the E-I pair’s coupling parameter (*C*). The activity of each phase neuron is defined by a system of differential equations, following previous work^6,9^, for E neurons:

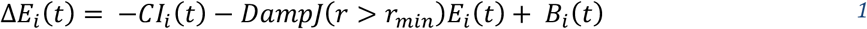

and for I neurons:

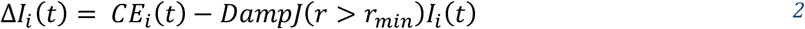

In which Δ*E_i_*(*t*) and Δ*E_i_*(*t*) denote changes in the E and I unit’s activity at each time step (model data were simulated at 500Hz, so Δ*t*=0.005s). The radius *r* of oscillation (*r* = *E*^2^ + *I*^2^) of an E-I pair is constrained to a radius *r_min_* = 1. To implement this constraint, we use an indicator function *J*, which returns 1 when *r < r_min_*, and 0 otherwise. The parameter *Damp* represents the strength of the damping, applied to constrain the oscillation’s radius to 1. *Damp* was set to 0.3 for processing nodes. For the MFC node, *Damp* was set to 0.005*theta_frequency to scale with the speed of the E-I pair theta oscillations and maintain an equal amplitude across time for all theta frequencies. The term *B_i_* denotes the burst that processing nodes could receive depending on the trial instructions (see **Medial Frontal Cortex unit** and **Lateral Frontal Cortex unit** for details). The MFC node did not receive bursts, thus *B_MFC_* was set to 0.

The frequency of oscillations generated by the E-I pair was defined by the coupling parameter *C*, and its relation to frequency in Hertz is given by the following equation:

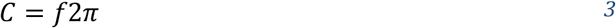

In which *f* denotes the frequency in Hertz and *C* the coupling parameter in the E-I pair.

Rate neurons receive, process, and transmit information to other nodes. Their activity (x_i_) is determined by inputs to the node (*in_i_*) and is modulated by its excitatory phase neuron (*E_i_*). Input to rate neurons of the Sensory unit, i.e. stimuli, were set at a value of 0.02. For instance, in a Sensory node, the input of the rate neuron is the intensity of the stimulus feature it is sensitive to (see **Processing units** for more details). The output *x_i_* of a rate neuron at time *t* is the product of *in_i_* and the activity of its excitatory neuron (*E_i_*(*t*)). Rate neurons activity are thus updated by:

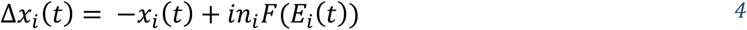

with *F*() being a logistic function of *E_i_*:

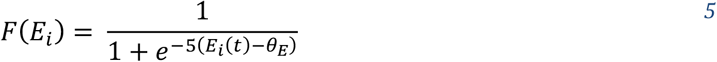

#### Processing units

The processing units are a Sensory unit and an Action unit. Each unit is composed of nodes representing cortical columns (see **Oscillatory nodes: a neuronal triplet** section).

In all nodes of Sensory and Action units, the coupling parameter *C* was set to generate gamma oscillations. The gamma oscillations were set to 30 Hz by using a coupling parameter of *C* = 188.5, which, in the computational implementation of the model, was set to 0.377 to account for the sampling rate at 500Hz (*C*/500 = 0.377). To show model stability and induce noise in processing nodes’ oscillatory phase, we introduced random slow fluctuations in the coupling parameter of nodes oscillating at gamma frequency. We generated random numbers from a normal distribution with parameters *μ* = 1, *σ* = 1, for each trial and each processing unit (i.e. Sensory and Action). A low-pass filter was then applied to these coupling fluctuations time courses, i.e. Gaussian convolution with *σ* = 1 (in seconds). Finally, the coupling parameter (i.e. *C* = 0.377, for 30Hz oscillations) was multiplied by the value of these low-frequency coupling fluctuations. The result of this manipulation was slow random fluctuations of gamma frequencies in phase neurons of processing units. For example, for one trial, Sensory nodes were oscillating at 32Hz at a certain time *t*, then gradually shifting to 27Hz, then to 35Hz, etc. This slow fluctuation was generated independently for Sensory and Action units.

Rate neurons of the Action unit receive input from rate neurons of Sensory nodes in order to implement the two-alternative orientation discrimination task on gratings. The main task was to report whether the target grating was tilted clock-wise (CW) or counter-CW (CCW) from the vertical axis. To report the tilt the rule was to use the index and middle fingers of either the left or right hand, indicated by the instructions. The left middle finger and right index finger should be used to report a grating tilted CW, and the left index finger and the right middle finger should be used to report a grating tilted CCW. Therefore, the connectivity between Sensory and Action nodes’ rate neurons implemented this rule.

#### Integrator unit

The Integrator unit accumulates information for each response, and triggers the model response once one of the Integrator nodes reaches a threshold. There is thus one Integrator node for each Action node. The Integrator nodes’ constitute a competitive accumulator network^11^ and followed the following update:

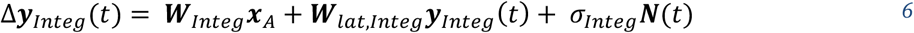

In which ***y**_integ_*(*t*) is a vector collecting the activity of all Integrator nodes at time t, ***W**_Integ_* denotes the weight matrix between Action nodes and Integrator nodes, ***x**_A_* denotes input from Action nodes to Integrator nodes. ***W**_lat,Integ_* denotes the update matrix of Integrator nodes in which off diagonal cells are set to −0.10 to implement lateral inhibition, and diagonal cells, representing the update rate of the competitive accumulator network, are set to 1. Finally, noise was added for each of the four variable Integrator nodes with *σ_Integ_* = 0.05 multiplying a vector *N*(*t*) of four random values drawn from a standard-normal Gaussian distribution.

As stated before, the Integrator unit produces a response when a threshold is reached by one of the Integrator nodes. To model a speeded task constraint, we implemented a collapsing threshold, equivalent to a collapsing bound in the drift diffusion model, which has been shown to adequately model the dynamics of response threshold in speeded tasks^12^. The threshold *θ_y_* therefore decreased exponentially from stimulus presentation to response deadline following this equation:

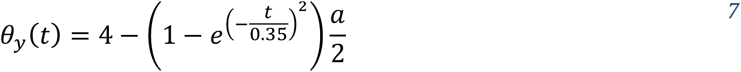

In which *θ_y_*(*t*) denotes the threshold of the Integrator unit at time t, and *a* denotes the initial starting point of *θ_y_*. In all simulations *a* was set to 4. Once one of the four Integrator nodes reached the threshold, we recorded the accuracy, depending on instruction, stimuli and the Integrator node which reach the threshold, and the time elapsed from stimuli onset, which provided reaction time for this response (see **Supplementary Fig.1c**).

#### Medial Frontal Cortex unit

The MFC unit generates theta oscillations that 1) generate bursts that phase-reset the processing units, and 2) initiate a competition window in LFC nodes.

The MFC unit is composed of one single node in which the E-I pair generates theta oscillations, whose frequency depends on the coupling parameter between the E-I pair. The rate neuron of the MFC node follows a Bernouilli process (Be) with a probability defined by the activity of the node’s E neuron:

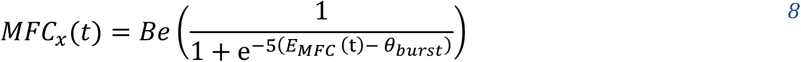

In which *Be* denotes the Bernoulli process, *E_MFC_* denotes the activity of the MFC E neuron, and *θ_burst_* (set to - 1) denotes the offset of the relation between *E_MFC_* and p (probability to trigger a burst). *Be*(*p*) is 1 with probability *p*, it will thus typically be 1 when *E_MFC_*(*t*) oscillation is near its peak. When *MFC_x_* = 1, a fixed amplitude burst = 0.5 is emitted to the LFC unit. The purpose of this burst is to synchronize processing nodes selected by the LFC, by phase reset of their E neuron (see **Processing units** for the burst effect and **Lateral Frontal Cortex unit** for the selection of the processing units receiving the burst).

In each cycle of theta oscillations in *E_MFC_* activity, a competition window is opened between rule nodes in the LFC. This competition window is opened when *E_MFC_ > θ_comp_*, with *θ_comp_* = 0.1. The competition window lasts a fixed temporal interval across cycles defined by a *θ_comp_* and the *C_MFC_* parameter, which determines the theta frequency.

To simulate different theta frequencies in the MFC, we varied the MFC coupling parameter (*C_MFC_*) from 0.050 (for 4Hz theta), to 0.087 (for 7Hz theta), see equation (3) in **Oscillatory nodes: a neuronal triplet**.

#### Lateral Frontal Cortex unit

The Lateral Frontal Cortex unit (LFC) contains rule nodes that act as pointers to processing nodes that constitute components of the rule. Rule nodes contain rate code neurons only, which together constitute a competitive accumulator network^11^. Each rule node receives a constant input throughout a trial from instruction nodes, which themselves are activated by the two instruction letters. Two instruction nodes represent the top letter of an instruction, and two other instruction nodes the bottom letter. This network of instruction nodes implements a congruency effect between instruction letters: top and bottom “Left” nodes were connected with a positive weight, thereby activating each other, and similarly for “Right” nodes (see instruction nodes in **Fig.1b** and **Supplementary Fig.1a**). Instructions are represented as a vector of binary values (zeros and ones) in which the first two indices represented a top L and R, respectively, and the two last indices represented the presence of a bottom L and R, respectively. For instance, the rule RL was represented as *instructions* = [0,1,1,0]. This was the input to the instruction nodes, which then projected to rule nodes through the following equation:

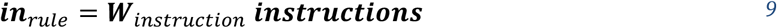

In which *in_rule_* denotes the input to rule nodes (i.e. from instruction nodes). Matrix ***W**_instruction_* represents the connectivity between instruction nodes implementing the lateral excitation, i.e. instruction letter congruency effect. The diagonal of ***W**_instruction_* was set to 1, and the cells representing the positive weight implementing the lateral excitation were set to 0.5.

The activity of rule nodes is updated through the following equation:

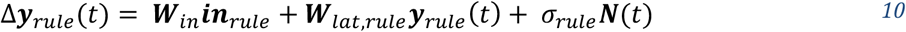

In which ***y**_rule_*(*t*) denotes the activity of all rule nodes at time t, ***in**_rule_* denotes the input to rule nodes (i.e. instructions) and ***W**_in_* denotes the weight matrix between instruction nodes and rule nodes in which weights between an instruction node of a particular letter and rules containing this letter was set at 0.5, e.g. instruction nodes “top R” and “bottom L” projected to the rule node “RL” with weight 0.5 (see connectivity between instruction and rule nodes in **Fig.1b** and **Supplementary Fig.1a**). ***W**_lat,rule_* denotes the update matrix of rule nodes in which off diagonal cells are set to −0.1 to implement lateral inhibition and the diagonal cells, representing the update rate of the competitive accumulator network, are set at 0.13. Finally, noise was added for each of the four rule nodes with *σ_rule_* = 0.075, multiplying a vector ***N***(*t*) of four random values drawn from a standard-normal Gaussian distribution.

The instruction nodes to rule nodes architecture allowed to manipulate task difficulty. For instance, the same-side rule LL, modeled as *instructions* = [1, 0, 1, 0], provided strong input to the LL rule node, and a small input to the LR and RL rule nodes as they each share the bottom and top letter, respectively, with the instruction LL. Thus, for *instructions* = [1,0,1,0], *in_rule_* = [0, 1.4, 0.7, 0.7], in which the *in_rule_* indices represent, in this order, RR, LL, LR and RL. On the other hand, a different-side rule like RL, modeled as *instructions* = [0, 1, 1, 0], provided a relatively strong input to the RL rule node, and a small input to LL, RR and LR nodes. Thus, for *instructions* = [0, 1, 1, 0], *in_rule_* = [0.7, 0.7, 0.4, 1], creating a stronger competition between the instructed rule (RL) and the other rules (RR, LL and LR).

Finally, the most activated rule node at each time *t*, amplified and routed the burst emitted at time *t* by the MFC (*MFC_x_*(*t*)) to the processing nodes it points to:

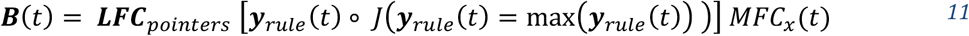

In which ***B***(*t*) is a vector of burst values arriving at each processing node’s E neuron resetting its phase. ***y**_rule_*(*t*) is the activity of rule nodes at time t and ◦ represents point-wise product. *JÇ)* is an indicator function that returns an array of 0 and 1, with 1 only for the most activated rule node at time t. ***LFC**_poirters_* is a matrix containing the processing nodes each rule node is pointing to. *MFC_x_*(*t*) is the activity of the MFC rate neuron at time t, i.e. this can be 0 or 0.5 (burst values were fixed) depending on whether the MFC is emitting a burst or not at that particular time point. Critically, only processing nodes composing the rule represented by the most activated rule node received the burst, while all other processing nodes did not. For instance, if the instructed rule is RR and the most activated rule node at time t is RR, the Sensory module “Right grating” and the Action module “Right hand” received the burst, thereby synchronizing their gamma oscillations.

As a result of the congruency in instruction letters and BC between rule nodes, the instructed rule will win the competition more often for same-letter rules, i.e. easy rules, than for different-letter rules, i.e. difficult rules (Supplementary Fig.2). Therefore, same-side rules will succeed to synchronize rule-relevant processing nodes more often. In the case of difficult rules, another consequence is that the model will achieve better performance at a slower theta oscillation frequency when competition lasts longer. In contrast, for easy rules, model performance increases with a slightly higher theta frequency. Hence, an optimal agent would shift theta frequency depending on task demands.

**Supplementary Figure 2.**
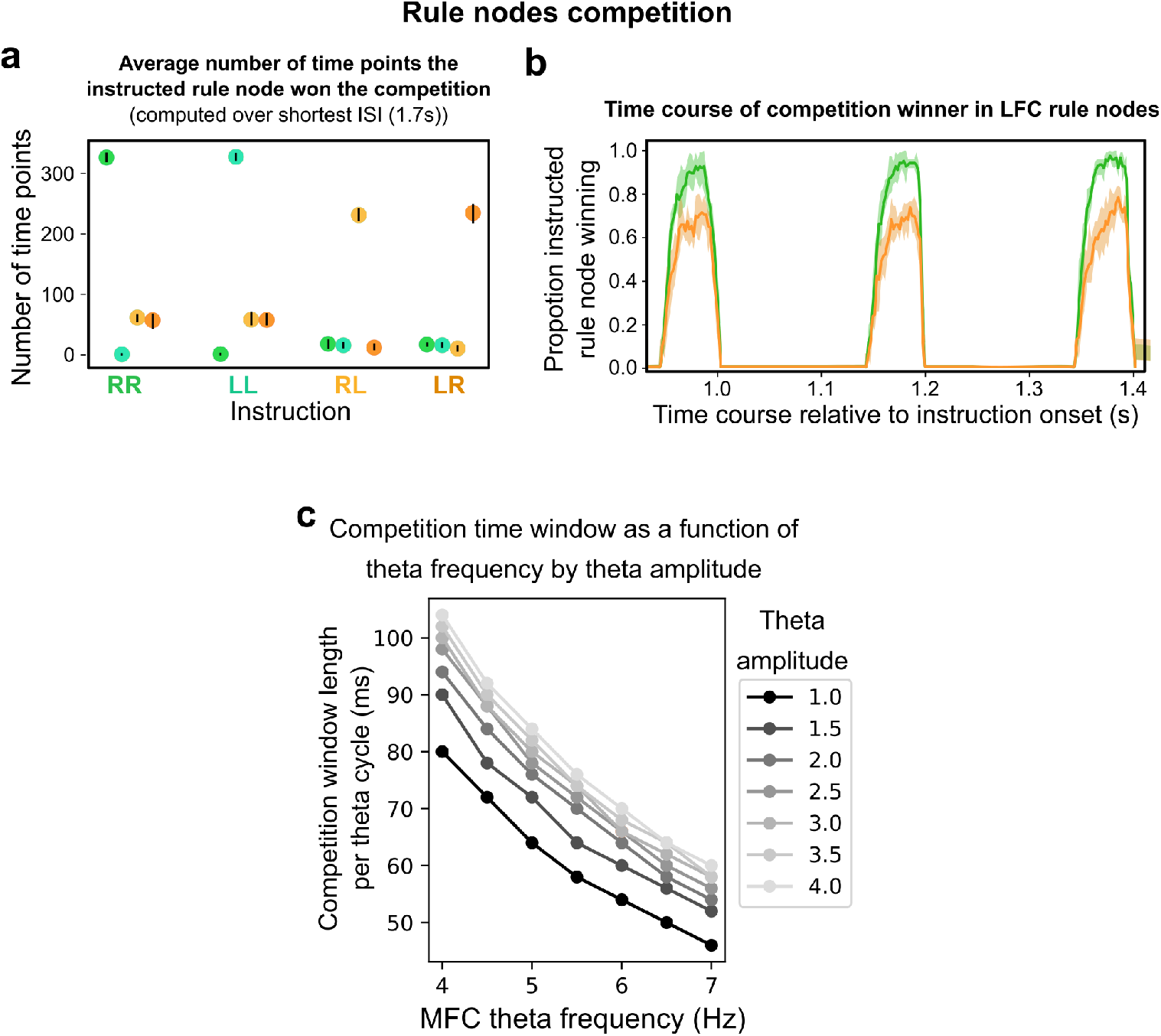
Rule node competition. **a.** Number of time points the correct rule node won the competition in the LFC unit depending on the instructed rule. Black bars represent the 95% Confidence Interval (smaller than dot size). **b.** Time course of proportion of correct rule node winning the competition across a sample of the ISD. Only two rule nodes are shown for clarity, one easy and one difficult. Shaded area represents the 95% Confidence Interval. **c.** Effect of MFC theta amplitude and frequency on competition time window. Average competition window length, in milliseconds, for one theta cycle, i.e. one competition window, as a function of theta frequency for different theta amplitude. Varying the amplitude (the different lines) shows that although the competition window increases with amplitude, this effect ceils around amplitude values of 2-3. Each line represents simulations at different MFC theta frequency with a fixed theta amplitude. Line color represents MFC theta amplitude as represented in the legend.

#### Simulations

We ran simulations of the model on the task depicted in **Fig.1a**. Instructions are shown for 200ms (two letters), then a variable ISD between 1,700 to 2,200ms, in 11 steps of 50ms, allows to prepare the instructed mapping, and subsequently two gratings are shown for 50ms. There were four possible instructions: RR, LL, LR and RL. There were four possible stimuli configurations: both gratings were tilted CW or CCW, or each grating was tilted differently than the other (e.g. left grating tilted CW, right grating tilted CCW). For each combination of task parameters, we ran 100 repetitions, which amounts to: (11 ISD + 4 instructions + 4 stimuli configurations) * 100 = 17600 trials. We then grouped repetitions into 34 groups of ~500 trials each, each representing one participant.

#### Effect of amplitude on competition window

To verify that high and low theta frequencies are optimal for easy and difficult tasks, respectively, we varied MFC theta amplitude and frequency and computed the competition windows length (**Supplementary Fig.2c**). Higher theta amplitudes increased the competition window length but quickly ceiled (around a value of 3).

Theta frequency on the other hand produced larger increases in competition window, indicating that effects of theta frequency on model task performance cannot be explained by theta amplitude alone.

**Supplementary Figure 3.**
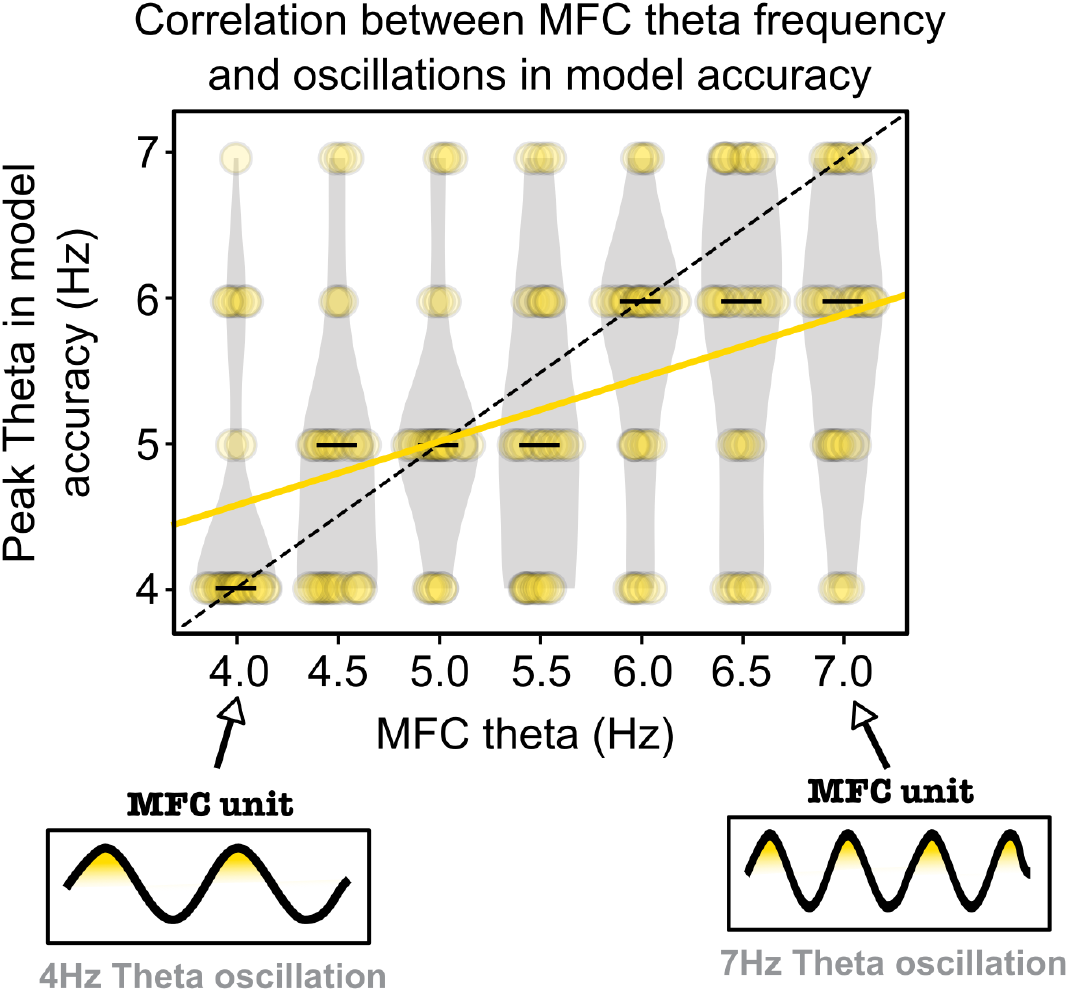
Correlation between MFC theta frequency and peak theta frequency in model accuracy across ISD. We simulated different MFC theta frequencies and estimated the peak theta frequency of oscillations in accuracy across ISD (pooled across all rules). The MFC theta frequency is represented on the x-axis and the estimated peak theta frequency of oscillations in model accuracy on the y-axis. Each violin plot represents the distribution of estimated peak theta frequency for multiple simulations at a fixed MFC theta frequency. Black bars in each violin plot represent the median of estimated frequency. The yellow line represents the linear regression of MFC theta frequency and estimated peak theta frequency. The correlation was highly significant (Spearman rank-correlation: *r*=0.42, *p*<10^−11^), showing that oscillations in model accuracy across ISD closely match with MFC theta frequency.

### Experiment

#### Participants

Thirty-nine human participants were recruited for this experiment (M ± STD = 23.7 ± 4.5 years old, range: 18-41 years old; 27 females). All participants had normal or corrected-to-normal vision and no history of neurological problems. All participants provided written informed consent and received monetary compensation for their participation. Five participants were excluded from the analysis because they completed less than 5 blocks, had less than 50 trials per instruction or due to poor behavioral performances (i.e. less than 50% overall accuracy). One participant was excluded because it was left handed. The experiment was approved by the local ethics committee (Faculty of Psychology and Educational Sciences, Ghent University). We recruited 35 participants to reach a total number of 30 or more participants after dropout, considering a ~15% drop-out rate due to noise-corrupted data or other problems related to participants’ task performance. The sample size after exclusions dropped at 29 participants, we thus tested 4 more participants, which were all included, bringing the sample size to 33 participants. Assuming a medium effect size and aiming for a power of 0.8 in a within-subject repeated measures ANOVA analysis, the study would require a sample of 32 participants, which makes our sample size sufficient and on par with current EEG research practices.

#### Apparatus and stimuli

Participants sat in a dimly lit room, 60 cm from a 24in LCD monitor (refresh rate: 60 Hz; resolution: 1280 × 1080 pixels). A chinrest was used to stabilize head position and distance from the screen. The experiment was implemented using Python 2.7 and the PsychoPy toolbox^13^.

#### Experimental design

Participants were instructed to perform a 2-alternative forced choice (2-AFC) orientation discrimination task on two sinusoidal gratings presented simultaneously on each side of a central fixation cross, as depicted in **Fig.1a**. Each grating was randomly tilted either CW or CCW relative to the vertical axis. The stimuli were sinusoidal gratings windowed by a raised cosine (size: 5° of visual angle, 10% contrast, 3 cycles per degree, at 5° eccentricity, on a gray background). The tilt angle was calculated for each participant using a staircase procedure (see below) to avoid ceiling accuracy. Participants were instructed at the beginning of every trial to perform the 2-AFC task on the right or left grating, and respond using their right or left hand (index and middle finger respectively for CW and CCW tilt).

Instructions letters were presented for 200ms with a size of 0.75° of visual angle, and positioned above and below the central fixation cross (vertical eccentricity: 1° of visual angle). The letter above the fixation cross instructed which grating was the target, i.e. on which grating the discrimination should be performed, and the letter below the fixation cross instructed which hand to use to respond. After instructions a preparation interval followed to allow participants to process instructions and prepare the stimulus-action mapping to perform the task. We used a dense behavioral sampling paradigm with multiple, densely distributed, instruction-stimulus delays (ISD)^14^: the duration of the ISD, between instructions and stimuli, was randomly chosen on each trial from 11 possible durations going from 1,700 to 2,200ms in 11 steps of 50ms. The variation in ISD was introduced to measure oscillations in behavioral performance and test predictions of the model (see **Fig.1f**).

A trial time course consisted of a 1000ms baseline period, followed by instruction presentation for 200ms. Then the variable ISD followed, which lasted between 1,700 and 2,200ms in 11 steps of 50ms, followed by stimuli presentation for 50ms. After stimuli onset, the fixation cross turned blue, indicating the beginning of the 700ms response window. If a correct response was given the fixation cross turned green, if an incorrect response was given the fixation cross turned red. If no response was given during the response window, a message indicated that the participant was too slow and paused the experiment, prompting the participant to take a break if needed, and press “Space” to resume the experiment. Every trial that was missed, i.e. not responded to, was added to the trial queue, and re-presented at the end of the block. Participants performed one training block to familiarize them with the experimental design, one staircase block to compute the participant’s grating tilt angle, and between 5 and 8 blocks of the task depending on the number of missed trials (i.e. participants who missed more response deadlines had longer blocks because trials were queued at the end of the block). The practice block consisted of 80 trials, the stimulus was shown for 100ms and the response window lasted 1000ms to make the task easier.

Following the practice block, participants completed a block implementing a staircase procedure on the tilts of the gratings. The staircase was done across all instructions and all ISDs to find a tilt level that would avoid ceiling performance and thus allow for fluctuations across ISDs. We used a one-up two-down staircase procedure consisting of 80 trials. The event timings and stimulus properties were the same as in the main task. Only the tilt of the gratings varied throughout the trials. Initially, a wide tilt (7°) was set. The procedure started with a step size of 3°, which was divided by 2 every second reversal starting at the second reversal. The reversal corresponded to switches in participants’ response accuracy, i.e. from a sequence of correct responses to an incorrect response or the other way around. When a participant switched from correct responses to an incorrect response, the difficulty of the task decreased by increasing the tilt of the gratings. Conversely, when a participant responded correctly after a sequence of errors, the difficulty of the task increased, i.e. decrease in the tilt of the gratings. The minimum tilt step size was set at 0.1°, the maximum final tilt of the gratings was 30° and the minimum was 0.5°. The final tilt was the average of the last 10 tilts.

**Supplementary Figure 4.**
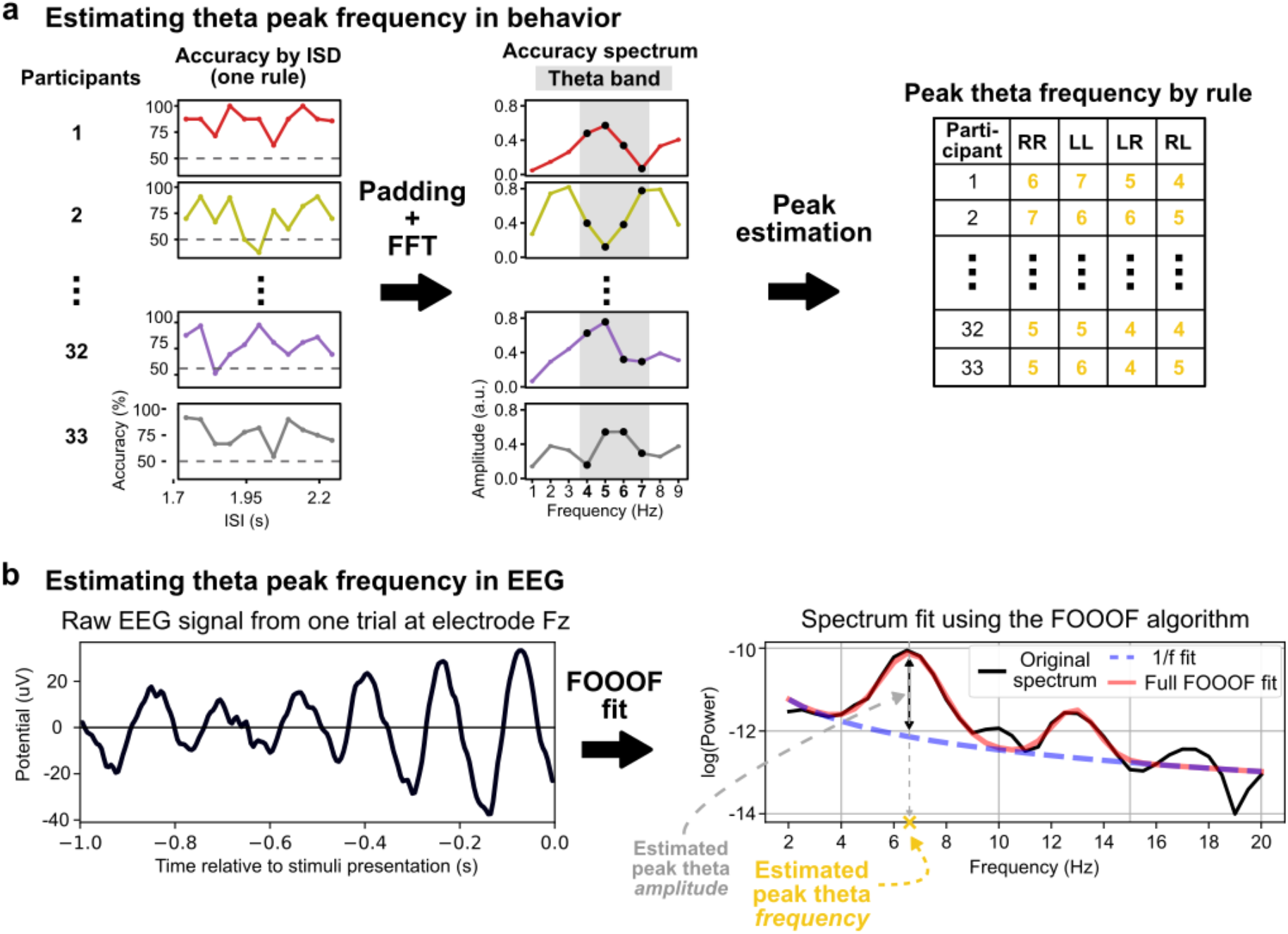
Theta peak estimation in behavior and EEG. **a.** Our model proposes that cognitive control relies on theta-rhythmic synchronization of task-relevant neural networks. This synchronization is done through bursts sent at the peak of MFC theta oscillations; therefore, around these phases (i.e. the peaks) of the theta oscillation, behavioral performance should be higher (yellow color in the oscillation’s background in Fig.1f). To test our model’s prediction that behavioral accuracy should oscillate at slower theta frequency for more difficult rules, we used densely distributed Instruction-Stimulus Delays (11 steps, 50ms apart) to sample behavioral performance at different phases of theta oscillations. To estimate peak theta frequency, we ran a Fourier Transform on padded participant accuracy-by-ISD, separately for each rule (only one rule shown in “Accuracy by ISD” and “Accuracy spectrum”). We then estimated the peak theta frequency (see Methods) in these behavioral accuracy spectra. This yielded a peak frequency (between 4 and 7Hz) per instruction per participant. **b.** To estimate peak theta frequency in EEG data we used a recent algorithm (FOOOF, see Methods) which parametrizes power spectra and extracts peak frequency and amplitude found over the 1/f pattern of the spectrum. The FOOOF algorithm first fits the 1/f pattern in the spectrum (purple dashed line), then estimates whether peaks over this 1/f pattern are present in the spectrum by fitting Gaussians. The parameters of these gaussians thus provides an amplitude and a peak frequency for the detected oscillation. Only one trial and one electrode (Fz) are shown in this figure for clarity. We then averaged the estimated peak theta frequencies using the FOOOF algorithm across the selected electrode cluster (see Methods, Fig.2c) for correct and incorrect trials separately. This yielded two peak theta frequency (one for correct trials, one for incorrect trials) for each rule for each participant.

After the staircase block, participants completed between 5 and 8 blocks of the main task depending on the number of trials missed, i.e. participants who missed the response deadline more often, had longer blocks (because of queued trials), and therefore completed less blocks. In total the experiment lasted ~3 hours from explanation of the task to removing the EEG cap.

#### Eye-tracking acquisition and processing

We recorded eye movements using a SMI eye tracker with a sampling rate of 250Hz (RED250 mobile system; SensoMotoric Instruments, Teltow, Germany). The eye-tracker camera using infrared optics was attached to the bottom of the computer screen. Each block of the experiment started with a calibration procedure in which participants had to follow a moving red dot with their eyes to nine locations on a grey background, the success of which was validated before continuing. Gaze position was epoched from instructions onset to stimulus presentation. To epoch gaze position data and align them with EEG data, we aligned the trial onset (instructions presentation) using the trial onset trigger in eye-tracking data and the trial onset trigger in EEG data. We then calculated the distance from the fixation cross in degrees of visual angle at each time point in the epoch. Any trial in which the gaze was outside a 1.5° radius centered on the fixation cross at any moment in the ISD, was rejected in the behavioral and EEG data.

#### Behavioral data analysis

As described above, trials in which gaze position’s distance from the fixation cross exceeded 1.5° of visual angle were discarded. Trials were grouped by instruction and by ISD. Model simulations showed a thetafrequency – rule-difficulty interaction in accuracy but not reaction times (**Fig.1d** and **Supplementary Fig.5a**). We therefore used accuracy as our dependent variable.

To compute spectra of behavioral accuracy oscillations across ISDs we first average-padded accuracy values (**Supplementary Fig.4b**). Average-padding was performed for each participant and for each instruction independently to increase frequency resolution to 1Hz. To pad the data, values corresponding to average accuracy across ISDs (by instructions) were added on either side of the empirical data points. Specifically, the 11 time points, spanning 500ms, were padded to get a 1,000ms segment, thus adding 4 data points before the first data point and 5 after the last one. Note that we also performed the analysis on nonpadded data and obtained similar results.

Then we computed a fast Fourier transform (FFT) to obtain frequency spectra of each accuracy time course across ISDs for each participant and each instruction. FFT allows to decompose the behavioral data from the time domain into frequency components to estimate an amplitude spectrum, i.e., the amount of each frequency present in the original data. To extract peak frequency in these spectra we computed the Laplacian transform of the spectra:

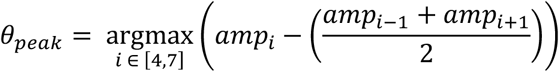

In which *amp_i_* denotes the amplitude at frequency *i*.

Finally, we centered the *θ_peak_* measures across instructions for each participant to discard any difference in offset of the *θ_peak_* measures across participants to test the model prediction that theta peak frequency in behavioral accuracy shifts depending on task demands.

**Supplementary Figure 5.**
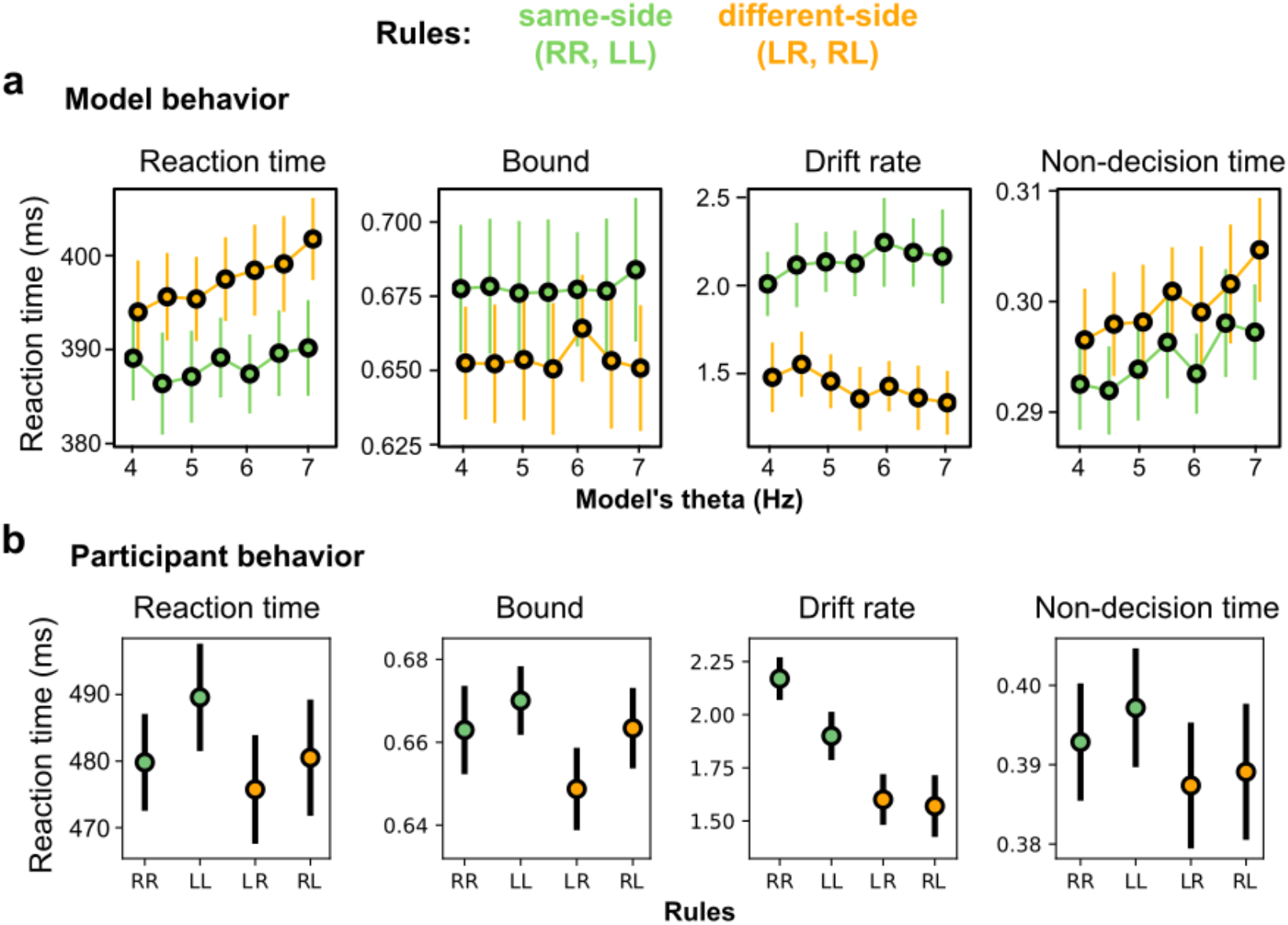
Reaction times and control analyses with DDM. Reaction times and DDM parameters (bound, drift rate and non-decision time) estimated in model and participant data with the EZ-Diffusion model (see Methods). **a.** Reaction time and DDM parameters of model performance by rule difficulty (same-side, different-side) and theta frequency (4 to 7Hz, steps of 0.5Hz). We ran a repeated-measure 2×7 ANOVA with factors rule difficulty (2 levels) and theta frequency (7 levels). There was a main effect of rule difficulty in all measures, i.e. reaction times, bound, drift rate and non-decision time, (all *F*(1,32)>116, all *p*<10^−10^). There was a main effect of theta frequency in reaction times, drift rate and nondecision time (all *F*(6,32)>2.6, all *p*<0.02). There was a significant rule-difficulty – theta-frequency interaction in reaction times (*F*(6,32)=3.6, *p*=0.0029) and drift rate (*F*(6,32)=7.8, *p*<10^−6^). Error bars represent standard deviation across simulations. **b.** Reaction time and DDM parameters (bound, drift rate and non-decision time) estimated on participants’ data grouped by rule. Data were collapsed across ISD to avoid data sparsity. We ran a repeated-measure 2×2 ANOVA with factors hand (2 levels: Left, Right) and target-location (2 levels: Left, Right). Only drift rate showed a significant interaction between hand and target-location (*F*(1,32)=26.36, *p*<0.00002), as well as a main effect of hand (*F*(1,32)=13.41, *p*=0.00089). All other effects were not significant (*F*(1,32)<3.8, *p*>0.05). Error bars represent s.e.m.

#### EEG acquisition and preprocessing

EEG was recorded using a Brain Products actiChamp system with 64 active scalp electrodes positioned according to the standard international 10–20 system at a sampling rate of 512 Hz. Four electrooculographic (EOG) channels were used to record eye-movements and blinks: two were placed on the outer canthi of the eyes, and two were placed above and below the right eye. All preprocessing steps were carried out with the Python MNE toolbox v.0.18.2^15^. Raw EEG data were downsampled offline to 200Hz, re-referenced to the average reference and low-pass filtered at 48Hz using a FIR filter with a Hamming window. The analysis of the pre-stimulus interval was performed on epochs from −1000ms to 0ms relative to stimulus onset, yielding epochs of 1000ms. A linear detrend was performed on each epoch individually. After trial rejection based on eye-tracking data (see **Eye-tracking acquisition and processing**) raw EEG and EOG time courses were visually inspected on a trial-by-trial basis to reject visible artifacts, eye movements or blinks. The average percentage of rejected trials across participants was 26% ± 14 (mean ± standard deviation).

#### EEG spectral analysis

To estimate peak frequency of theta oscillations we first computed power spectral density over the 1000ms window using Welch’s method provided in the Scipy toolbox v.1.3.1^16^. The Welch power density estimation was performed using a Hann window and zero-padding to obtain 400 time points of data in order to smoothen the spectra, as this improves estimation of peak frequency in the following analysis step. We then used a recent method that allow to parametrize neural spectra by fitting the 1/f pattern in electrophysiological recordings spectra and providing a sensitive identification and estimation of oscillatory processes in neural activity (FOOOF toolbox, version 1.0.0^17^). This algorithm yields several measures, including the peak frequency and amplitude of oscillations detected over the 1/f pattern in the spectra (see **Supplementary Fig.4c**). Using this algorithm, we computed separately for every participant, trial and electrode, whether a peak was detected in the theta frequency range (i.e. higher than 3Hz, and lower than 8Hz) and saved the estimated peak (in Hertz) and the amplitude of the peak (in μV^2^/Hz). Settings for the FOOOF algorithm were set as follows. The power spectra were parameterized across the frequency range 2 to 20Hz. The peak width limits were set between 0.5 and 2, to find peaks that were frequency specific. The maximum number of peaks was set at 4, under the assumption that in the 2-20Hz frequency range there could be four meaningful peaks, i.e. one in each band (delta, theta, alpha and beta). No minimum peak height was set, peak threshold was set at 2 (default), and aperiodic mode was fixed (default).

To test model predictions in theta peak frequency we separated trials according to the instruction and accuracy for each participant and each electrode. As a sanity check, replication of previous findings on theta amplitude, and as an independent electrode selection procedure, we investigated the scalp distribution of theta oscillation amplitude for correct versus incorrect trials. In order to perform this analysis, we z-scored the amplitude values extracted using the FOOOF toolbox, across all electrodes for each instruction, for each trial separately. This allowed to highlight the specific theta amplitude topography elicited by proactive cognitive control, i.e. preparing to implement an instructed stimulus-action mapping. We performed a cluster-based permutation test^18^ with 10,000 permutations on scalp topographies to test whether a cluster of electrodes showed relatively higher theta amplitudes in correct versus incorrect trials (across all instructions). This analysis revealed a significant cluster of electrodes in fronto-central sites (p=0.0001, **Fig.2c**).

Finally, to test the model prediction about shift in optimal theta frequency across task difficulty, we computed the average peak theta frequency (in Hertz), extracted using the FOOOF toolbox, in the selected cluster of electrodes (**Fig.2c**). We z-scored the peak frequency value across rules, separately for each participant, to discard any difference in offset or range of the EEG theta peak frequencies across participants. This procedure was carried out to specifically test the model prediction that theta peak frequency decreases with task difficulty, thus inter-individual differences in theta peak frequency for each instruction were not of interest in this specific analysis.

#### Statistical analyses

To compute the optimal theta frequency per rule difficulty (same-side (RR, LL) versus different-side (LR, RL) rules) in the model we calculated the MFC theta frequency yielding the highest accuracy for each group of simulations (see **Simulations**). We then compared the two samples of optimal theta frequencies per rule difficulty using a two-sided Student t-test (**Fig.1e**). For all correlations we used a Spearman rank-correlation (**Fig.2e** and **Supplementary Fig.3**).

Reaction times and DDM parameters estimated on model data (**Supplementary Fig.5a**) were analyzed using a 2-by-7 repeated measure ANOVA with factors rule difficulty (two levels: same-side and different-side) and theta frequency (seven levels: from 4 to 7Hz in steps of 0.5Hz), using the Python StatsModels package v0.10.1 (https://www.statsmodels.org/v0.10.1/).

Participants’ behavioral and EEG data were entered into a two-way repeated measure ANOVA with factors hand (two levels: Left, Right) and target-location (two levels: Left, Right) using the Python StatsModels package. For behavioral data it consisted of accuracies, reaction times, and DDM parameters per rule (**Fig.2a; Supplementary Fig.5b**), and peak theta frequency (i.e. of accuracy-by-ISD) per rule. For EEG data it consisted of the average peak theta frequency from the selected electrode cluster (Methods) of correct trials per rule (**Fig.2d**; circles); and the difference between average peak theta frequency of correct and incorrect trials.

To investigate inter-individual differences in the sensitivity of EEG peak theta frequency to rule difficulty we performed a linear regression of each participant’s EEG peak frequency (correct trials) ordered by each rule’s overall accuracy across participants (i.e. rule was treated as a linear predictor: RR=76.27, LL=72.44, LR=69.24, RL=68.08). In a second step, individual-participant slopes were correlated with overall accuracy, collapsed across rules (**Fig.2e**).

#### Data and code availability

Raw behavioral, eye-tracking and EEG data can be found at this Open Science Framework repository: https://osf.io/nwh87/?view_only=b11ee1f860804da582c816fe8acdecad

Code of the model, the behavioral experiment and analysis scripts to reproduce all results and figures from the study can be found on this Github repository: https://github.com/mehdisenoussi/theta_shift_cog_control

